# Can Stephen Curry really know? - Conscious access to outcome prediction of motor actions

**DOI:** 10.1101/2021.03.30.437477

**Authors:** Lisa K. Maurer, Heiko Maurer, Mathias Hegele, Hermann Müller

## Abstract

The NBA player Stephen Curry has a habit of turning away from the basket right after taking three-point shots, presumably because he can predict the success of his shot. For such a consciously accessible prediction to be possible, Stephen Curry needs access to internal processes of outcome prediction and valuation. Computational simulations and empirical data suggest that the quality of internal prediction processes is related to motor expertise. Whether the results of internal predictions can reliably be consciously accessed is less clear. In the current study, 30 participants each practiced a virtual goal-oriented throwing task for 1000 trials. Every second trial, they were required to verbally predict the success of the current throw. Results showed that on average, conscious prediction accuracy was above an individually computed chance level, taking into account individual success rates and response strategies. Furthermore, prediction accuracy was related to throwing performance. Participants with better performance predicted the success of their throws more accurately than participants with poorer performance. Moreover, for the poorer performing individuals, movement execution was negatively affected by the verbalized predictions required, and they did not show variation in speech characteristics (response latency) between correct and incorrect predictions. This indicates reduced quality of conscious access to internal processes of outcome prediction.

## 1 Introduction

Theories of internal models claim that the predictive outcomes of forward models play an essential role in motor learning (McNamee & Wolpert, 2019). Forward models are a set of neural processes that integrate information from the current state of the system and its environment with motor commands and sensory signals from movement execution to predict sensory consequences of that movement (Miall & Wolpert, 1996). Sensory and motor noise increase the uncertainty in the state estimate of the system. Forward models act as filters capable of reducing this uncertainty and attenuating unwanted information, or highlighting information critical for control. Furthermore, forward models can be used to transform motor errors, which are differences between desired and actual sensory outcomes of a movement into corresponding corrections in motor commands, thereby providing appropriate signals for motor learning (Wolpert et al., 1995). Based on computational simulations, it is suggested that learning is faster the better the forward model is able to model the dynamics of the movement and its effects (Jordan & Rumelhart, 1992). In line with this, empirical and anecdotal evidence confirm that processing and valuation of motor errors based on forward model predictions is strongly related to learning. First, it has been shown that neurophysiological correlates of predictive error valuation increase with learning (Beaulieu et al., 2014; Lutz et al., 2013; Maurer et al., 2021). Second, participants with extended experience in a throwing task particularly motor experts, that is throwers with high accuracy show more distinct signs of predictive error valuation on the neurophysiological level (Joch et al., 2017; Maurer et al., 2015; Maurer et al., 2021). Third, there is anecdotal evidence that motor experts are aware of their own errors and can consciously predict them even before they perceive any external feedback: The alleged predictive abilities of NBA players like Stephen Curry can be observed in both professional and amateur game videos, and in them it can be observed that shortly after the ball leaves the player’s hand, the players already cheer in cases of successful throws or express their disappointment in cases of missed throws. But is it really possible to gain conscious access to the predictive output of forward models in highly complex motor tasks like basketball shooting? And taken even further, might conscious access to prediction processes influence forward model computations? If so, would this influence be beneficial, as it has been shown in an apparent motion paradigm (Vetter et al., 2014), or detrimental, as proposed by the theory of reinvestment (Masters et al., 1993; Masters & Maxwell, 2008), and indicated by studies on perception-action coupling (Beilock et al., 2002; Farrow & Abernethy, 2003)? Without delving deeply into theories and models of consciousness, it has to be acknowledged that the term “consciousness” has different meanings, such as reaching from a waking state or subjectivity to an experimental variable of brain differences attributable to consciousness (Baars, 2015). In the present study “conscious access” is defined as the ability to report contents of perceptual states; according to this definition, perception itself is not limited to the processing of sensory (afferent) signals, as perception can arise from afferent and efferent information as well as cognitive (top-down) signals. Hence, if people with reliably working forward models have conscious access to the output of the forward model, they should be able to verbally predict the outcome of a motor action before any external feedback about the outcome is available. Arbuzova and colleagues (2021) examined metacognitive abilities in the discrimination of two different outcomes based on predictions in a virtual goal-oriented throwing task. Discrimination accuracy was governed by an online staircase procedure aimed at fixing performance at approximately 71% correct. Results from this study do not allow conclusions about absolute discrimination accuracy (i.e., effect prediction), but, confidence about the discrimination ratings (metacognitive ability) was relatively high across different informational domains (visual, visuomotor, motor). This shows that aspects of one’s own movement execution are principally available for conscious use, at least with respect to the monitoring of performance.

The conscious accessibility of sensorimotor prediction with respect to action outcomes has been investigated in athletes involved in different sports. The focus of most studies has been on anticipatory estimates of other players’ actions, based purely on visual information, for example in basketball shooting (e.g., Abreu et al., 2012; e.g., Aglioti et al., 2008; Li & Feng, 2020; Özkan et al., 2019), volleyball smashes (e.g., Cañal-Bruland et al., 2011; Wright et al., 1990), soccer penalty kicks (Tomeo et al., 2013), or other game situations (for a review see Abreu et al., 2017). Since predictions in these studies were exclusively based on observations of motor actions, available sensorimotor information was incomplete, and visual information differed compared to when actions were executed. That is, observers lacked internal efferent information about motor commands, and did not have access to proprioceptive or haptic information associated with the related movement either. But, in contrast to performers, observers take a third-person perspective. Hence, they can use visual information from whole body kinematics. Other studies examined predictions in both observers and performers (Cañal-Bruland et al., 2015), or in performers alone (Maglott et al., 2019). Performers had to rate outcomes of basketball shots after their vision was occluded by shutter googles immediately after ball release. Results showed that, on average, performers were able to verbally predict the results of their shots above the level of chance. But, both of these studies reported a strong judgment bias regarding shooting position and outcome (hits vs. misses). Performances of shots from the foul line were generally overestimated as compared to shots taken from other distances (Canal-Bruland et al., 2015), and expert players in particular had higher biases towards predicting their shots as hits (Maglott et al., 2019). The authors took that bias into account when calculating the base rate of correct judgements (i.e., the number of all actual hits being predicted as hits plus the number of all misses being predicted as misses, relative to the total number of trials) that could be accounted for by pure chance. Yet, the base rate reflecting pure chance also depends on the actual individual hit rate, which, however, was not included in the base rate estimations provided by the authors. Furthermore, in both studies, the shutter googles were manually controlled, which presumably introduced relatively large temporal variations of the occlusions, and it must be assumed that some post-release information about ball trajectory could have been received and processed by participants.

The present study aimed to verify that subjects with experience in a motor task can consciously predict outcomes of their own actions without any external feedback about action outcomes being available to them. For the experimental task, a virtual goal-oriented throwing task with parallels to basketball shooting was used. One significant advantage of studying throwing in this context is the natural delay between movement (throwing) termination and the availability of outcome feedback. Moreover, since the task was virtual and movement execution was captured online, the visual information available to subjects could be precisely controlled. Thus, outcome predictions could be based exclusively on information gathered during movement planning (efferent information) and during movement execution (haptic, proprioceptive, or visual information), but not on external information about movement effects (e.g., trajectory of the object to be thrown). Hence, information on the level of an individual’s accuracy in consciously accessing outcome predictions (predictive accuracy) would provide novel insight into the quality of forward models and the easiness or efficiency of conscious access to forward models. Predictive accuracy was quantified by the rate of correct verbal predictions of throwing outcome, relative to a baseline (chance) level accounting for individual hit rates and response strategies.

Successful verbalized predictions require at least two separate functions: a predictor and conscious access to its predictions. Or, conversely, poor predictive accuracy may arise from two reasons: (i) individuals have poor prediction quality (due to poor forward models), or (ii) they have difficult or inefficient conscious access to their forward models. Thus, it is expected that individuals with superior throwing performances (inferring good forward models) and easy conscious access to internal processes, would be able to predict their throwing outcomes above the level of chance. In the present study design, the integrated effect of both aspects was examined, and experimental separation was not directly possible. But, post-hoc interpretations of the different influences of prediction quality (i) and easiness of access (ii) on prediction accuracy are provided. As additional variables contributing to this differentiation, throwing performance and speech characteristics of verbal responses were analyzed. A possible back wash effect of conscious access to internal prediction processes results in interference with throwing performance: The preparation of conscious verbal predictions following movement execution might disrupt the motor control process, and hence affect throwing performance (Beilock et al., 2002; Masters & Maxwell, 2008). These costs of conscious processing might also manifest themselves in longer verbal response latencies and lower response volumes due to hesitation (Collins et al., 2000; Seymour, 1970).

## 2 Materials and Methods

### 2.1 Participants

Thirty participants (18 female, 12 male) from the student population of the Justus Liebig University, Giessen, Germany with an average age of 24.13 (*SD* = 5.77) years participated in the study. Participants were healthy and had normal or corrected-to-normal vision. Two left-handed subjects practiced the task with the right hand, which had been shown to produce similar learning curves to right-handed participants in pilot studies. Participants received course credit or monetary compensation of €8 per hour. The experiment was conducted in accordance with the ethical standards laid down in the Declaration of Helsinki. The protocol was approved by the Ethical Review Board of the Justus Liebig University, Giessen.

### 2.2 Experimental task and apparatus

Participants practiced a novel and complex goal-oriented throwing task that has previously been used to study motor learning (e.g., Cohen & Sternad, 2009; Maurer et al., 2021; Müller & Sternad, 2004; Pendt et al., 2011). The task is inspired by the British pub game “Skittles”, where a ball attached to the top of a post by a string has to be swung around the post to hit target objects on the opposite side. In addition to the ballistic nature of the task preventing online corrections during movement execution, this throwing task allows a temporal separation of movement execution and its terminal outcome, because the outcome is temporally delayed with respect to the movement. The task was executed semi-virtually. That is, participants executed a real ballistic throwing movement using a metal lever device (manipulandum), while the movement and its outcome were only visible on a computer screen from an overhead perspective (see Fig. 1).

**Figure 1.**
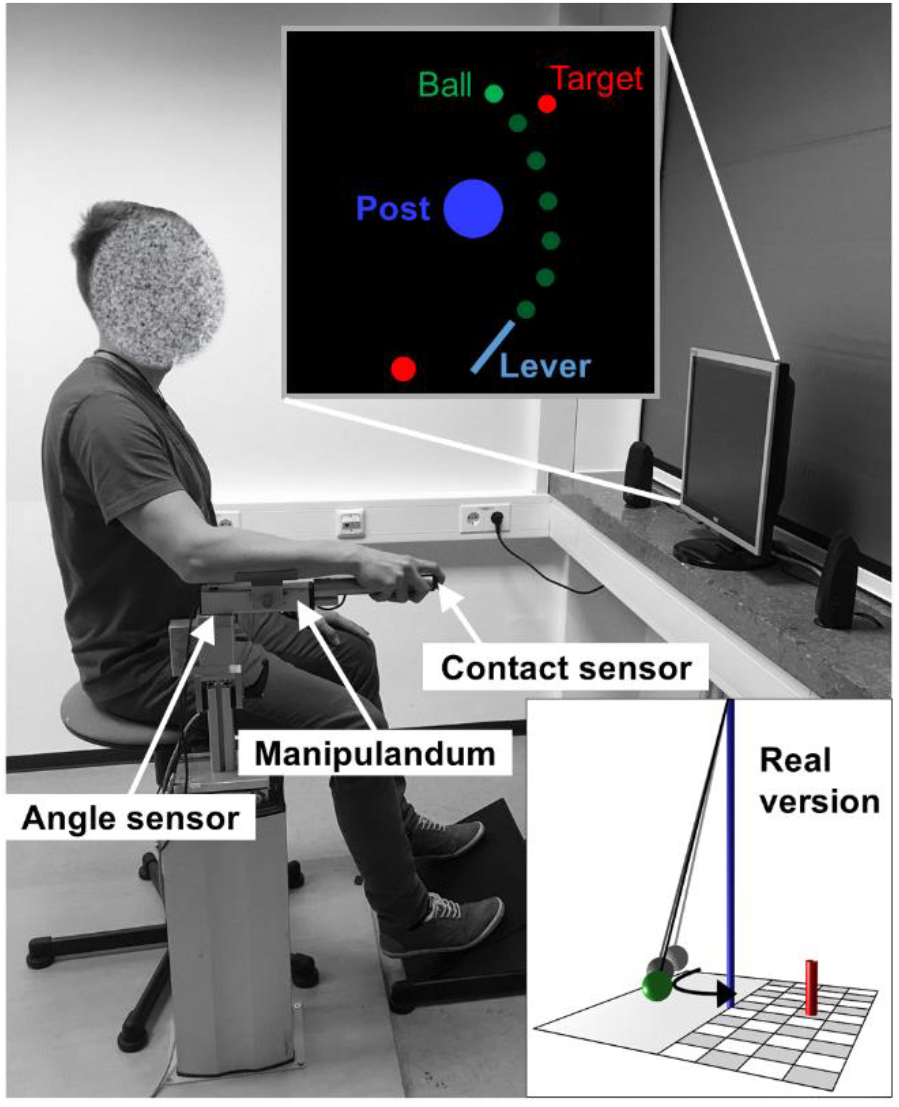
Experimental setup of the Skittles task. The participant uses a manipulandum to throw the virtual ball (green) with a horizontal rotational movement. The ball is released by lifting off the index finger from a contact sensor at the tip of the manipulandum. The ball travels on an elliptical pathway around the blue post to hit a red target.

The Skittles task was carried out using MATLAB R2018a (The Mathworks, Inc.) using the Psychophysics Toolbox version 3.0.14 (Brainard, 1997). On the screen in front of each subject, a virtual equivalent of the metal lever was displayed, which participants used to pick up and throw a green virtual ball (radius on screen = 4.2 mm) around a blue center post (radius on screen = 21 mm) in order to hit a red target (radius on screen = 4.2 mm). The elliptical trajectory of the ball around the center post was defined by the angle and velocity of the manipulandum at the moment of ball release. The calculation of the ball trajectory was based on a physical model of the task (Müller & Sternad, 2004) with the following parameters: center post (radius = 0.25 m; position: x = 0.0 m, y = 0.0 m), target (radius = 0.05 m; position: x = 0.8 m, y = 0.9 m), ball (radius = 0.05 m; mass = 0.1 kg), spring constant (1.0 N/m). In the regular version of the task, participants were able to see the ball moving towards the target after ball release. Whenever the minimum distance between the trajectory of the ball center and the center of the target (D_min_) was less than or equal to twice the radius of the ball/target, the ball collided with the target (hit). For the subjects, this was apparent visually, because the target was pushed away from its position, and acoustically by the sound of two colliding billiard balls. In trials where D_min_ was larger than twice the radius of the ball/target, the ball missed the target. In the experimental version, in every second trial the ball vanished from the screen immediately after its release from the virtual lever. In these trials, subjects did not receive any information about the movement outcome.

### 2.3 Study procedure

Task execution was accomplished as follows: Participants sat on a stool placed 100 cm in front of a 19-inch, 4:3 computer monitor (model: Dell P190St, screen resolution: 1280 x 1024 pixels, refresh rate: 60 Hz). Their right arms rested on the foam padded manipulandum, which was fixed on a height-adjustable stand at the vertical rotation axis below the elbow joint of the participant (Figure 1). An integrated magnetic angle sensor with a resolution of 12 bit (0.09 deg) measured the lever rotation with a sampling rate of 1000 Hz. Movement was restricted to the horizontal plane, more specifically, to rotation around a fixed vertical axis. To pick-up the virtual ball, participants placed their index fingers on an electrical contact sensor at the tip of the lever to close an electrical circuit. They then “threw” the ball by moving the manipulandum in an outward horizontal movement similar to a Frisbee toss, and starting in front of their bodies. As soon as the participant’s finger was lifted from the manipulandum, the virtual ball was released from the virtual lever. To explain the task to the participants, a miniature model of the real Skittles game was used to clarify the task. To prevent a fast, rhythmic execution of subsequent trials, participants were instructed to start every trial by moving the tip of the virtual lever into a red circle positioned left of the fixed end of the lever (35° clockwise relative to the horizontal axis; see Figure 1). Immediately after the tip of the virtual lever reached the circle, it turned yellow. The circle turned green when the lever was held steady within the yellow circle for one second. The green starting circle signaled that participants were free to start the movement at any time. Note, however, that the subjects did not start the movement in reaction to the green signal. The aim of the task was to hit the target as frequently as possible.

The movement result was to be predicted verbally by the subjects within 2.5 seconds after ball release by the German words for hit (“Treffer”) or miss (“Fehler”). The verbal utterances were recorded for later analysis. For this purpose, a clip-on microphone (Monacor ECM-501L/SK), a phantom power adapter (MG STAGELINE EMA-1), and a microphone preamplifier (IMG STAGELINE MPA-202) were used. The output signal from the preamplifier was captured using a 16-bit data acquisition device (National Instruments PCIe-6321), and Matlab Data Acquisition Toolbox time synchronized with the data from the Skittles apparatus (angular and touch sensor) with a sampling frequency of 10.000 Hz.

### 2.4 Study design

Practice took place over two sessions with 500 trials each. Trials were categorized with respect to practice and experimental conditions (overview in Tab. 1). The first 100 trials of session one served as a first practice of the task. The regular version of the task was used for all of these trials. From trial 101 to trial 150, the experimental version was used where ball flight information was masked for every second trial as described above (“Experimental task and apparatus”). In these trials, participants did not receive any outcome feedback, while feedback was normally presented in the other 50 % of trials. From trial 151 to trial 1000, the regular version of the task was alternated with the experimental version in every other trial and, additionally, participants were asked to verbally predict the outcomes of their throws in the trials without ball flight information and outcome feedback (*prediction condition*). Trials 151-200 served as practice of the verbal prediction. Only trials 201 to 1000 were used for analyses where the *prediction condition* (verbal prediction and no outcome feedback) was contrasted with the *regular condition* (no verbal prediction, but available outcome feedback). The alternation of throws with and without feedback was chosen because pilot data indicated that it was not possible to perform the task successfully without “drifting away” from the solution manifold of the task without regular feedback.

**Tab. 1.** Overview of the experimental procedure.

|  | **Trials 1-100** | **Trials 101-150** | **Trials 151-200** | **Trials 201-1000** |
| --- | --- | --- | --- | --- |
| **Experimental phases** | <u>Practice:</u><br>Regular task<br>version | <u>Practice:</u><br>Alternation of<br>regular task | <u>Practice:</u><br>Alternation of<br>regular task | <u>Experiment:</u><br>Alternation of<br>regular task |
|  |  | version and<br>experimental<br>version in every<br>trial | version and<br>prediction<br>condition in<br>every trial | version and<br>prediction<br>condition in<br>every trial |
| **Implementation**<br>**of feedback and**<br>**verbal**<br>**prediction in**<br>**the different**<br>**phases** | 100 % of trials<br>with outcome<br>feedback | 50 % of trials<br>with outcome<br>feedback and 50<br>% of trials<br>without<br>outcome<br>feedback | 50 % of trials<br>with outcome<br>feedback and<br>50 % of trials<br>without<br>outcome<br>feedback and<br>with verbal<br>prediction | 50 % of trials<br>with outcome<br>feedback and<br>50 % of trials<br>without<br>outcome<br>feedback and<br>with verbal<br>prediction |

### 2.5 Analysis of throwing performance

Behavioral analyses as well as the analyses of verbal responses were done in MATLAB R2020b (The Mathworks, Inc.). The execution variables (release angle and velocity) and outcomes in the Skittles task are related in a nonlinear fashion. Furthermore, the task is redundant, what means that hits and misses are not functions of a dichotomous difference in throwing execution, but can arise from very different combinations in release angle and velocity. As a consequence, outcome prediction is not trivial. To account for this difficulty, only clear target hits (with D_min_ ≤ 7 cm) and clear misses (with D_min_ ≥ 12 cm) were analyzed (following Joch et al., 2017; note that trials with Dmin ≤ 10 cm lead to hits). Throwing performance was defined as the rate of clear hits in percent of blocks of 100 trials (10 blocks in total) averaged over all participants. Since conscious verbal predictions could influence throwing performance, hit rates between the *prediction condition* and the *regular condition* were compared. In these cases, hit rate was determined over 50 trials of each block for both conditions, because prediction trials and regular trials were alternated every trial.

### 2.6 Analysis of verbal responses

Verbal predictions of throwing outcomes were examined along two dimensions: First, prediction accuracy was analyzed in order to test whether participants were able to consciously access their internal processing of movement errors. Second, characteristics of speech, concretely variance in the onset and amplitude of verbal responses provided further information about the ease of conscious access to the predictions. It was assumed that faster (easier) conscious access would be manifested in earlier and louder prediction responses.

#### 2.6.1 Prediction accuracy

Prediction accuracy was defined as the rate of correct predictions relative to an individual baseline or chance level. This was accomplished in several steps. First, the rate of correct predictions (%C_Pred_) was defined as the percentage of correctly predicted clear hits and misses of all trials in the *prediction condition*. This empirical prediction rate was compared to the individually calculated prediction baselines (%C_Chance_). This baseline depends on two factors: first, the rate at which participants actually hit (%Act_Hit_) or missed (%Act_Miss_) the target in the experimental condition and, second, the rate at which they verbally report hits (%Verb_Hit_) or misses (%Verb_Miss_).

Based on the null hypothesis that verbal estimates are unrelated to the actual occurrence of hits and misses, we estimated %C_Chance_ in the following way:

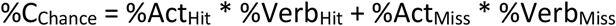

Prediction accuracy (%Acc_Pred_) was then computed as the percentage of correct predictions above %C_Chance_, normalized with respect to perfect predictions:

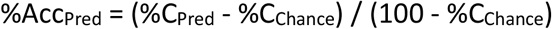

Thus, %Acc_Pred_ represented the ability of participants to consciously and verbally predict the outcomes of their throwing movements.

#### 2.6.2 Speech characteristics

To analyze verbal utterances, voltage values from the microphone output were offset-corrected, rectified, and then a moving average calculation (window width 250 values, i.e. 25 ms) was performed. The onset time of a verbal utterance was identified when the averaged profile exceeded 0.1 V. To determine the amplitudes, the maximum value in the averaged voltage curve was first determined. To account for general differences in loudness resulting from different positioning of the microphone and speaking volumes of subjects, all maximum amplitude values of each test session and subject were divided by their medians. Finally, the medians of onset latencies and response amplitudes were determined for all trials in the four different categories Act_Hit_/Verb_Hit_, Act_Hit_/Verb_Miss_, Act_Miss_/Verb_Hit_, Act_Miss_/Verb_Miss_.

### 2.7 Statistical analyses

Statistical analysis was performed in JASP (Version 0.14.1). The alpha level was set to .05 for all statistical analyses. Data normality was checked with the Shapiro-Wilk test, and sphericity was checked using the Mauchly’s W test. In case of violation of sphericity, the Greenhouse-Geisser correction was used. Changes in performance were tested with repeated measurement ANOVAs over all sessions, which included *Holm* corrected post-hoc testing of single sessions. Direct comparisons of performances between the experimental conditions (*regular* vs. *prediction*) was done using Wilcoxon signed-rank test (due to violation of normality assumptions), using the rank-biseral correlation coefficient as effect size. A one-sample *t*-test was used to examine whether prediction accuracy (%Acc_Pred_) was above baseline prediction, with effect sizes determined by Cohen’s *d*. It was expected that prediction accuracy would be a function of forward-model quality, and quality of conscious access. Hence, a Spearman correlation between prediction accuracy and hit rate as well as between prediction accuracy and the differences in hit rates between the *regular condition* and the *prediction condition* (conscious processing costs) was conducted. Speech characteristics (response latency and amplitude) were tested by a 2 (actual hit or miss) × 2 (verbalized hit or miss) repeated measures ANOVA, with the difference in hit rate between the *regular condition* and the *prediction condition* as a covariate. In addition, a Bayesian inference approach was used, with Bayes factors (*BF*) interpreted as the amount of evidence for the null and the alternative-hypothesis before versus after inspection of the data (Verdinelli & Wasserman, 1995). The size of the *BFs* were interpreted according to Raftery (1995). *BFs* 1 - 3 were interpreted as weak evidence for the alternative hypothesis against the null hypothesis, 3 - 20 as positive, 20 - 150 as strong, and *BFs* > 150 as very strong evidence.

## 3 Results

### 3.1 Throwing performance

Figure 2 depicts the development of throwing performance (hit rate). Figure 2A shows the hit rate over all trials executed. In Figure 2B, only trials carried out under regular conditions are shown (from trial 201 on), and Figure 2C illustrates the hit rates of only those trials of the *prediction condition* (from trial 201 on). Hit rates started at around 50 % on average, and rose with practice until they levelled off around block seven. ANOVA analyses carried out with repeated measures showed a significant main effect of block (*F*(4.65, 134.85) = 18.92, *p* < .001, *η*_*p*_^*2*^ = .40, *BF*_*10*_ > 150). Post hoc tests revealed significant differences between blocks 1-6 and block 10, practically no differences between block 7 and 10, and no differences between blocks 8, 9 and 10 (see Tab. 2). Interindividual variance was relatively large in general (standard deviation of the hit rate of all trials was 13.49 %, see Tab. 3), and even larger in the *prediction condition* (SD = 22.71 %). The average hit rate also differed significantly between the *regular condition* and the *prediction condition* (*W* = 61, *p* < .001, *rank-biserial correlation* = -0.74, *BF*_*10*_ > 150), but there was also large variance between participants (SD = 20.73 %, Min = -5.24 %, Max = 60.33 %).

**Figure 2.**
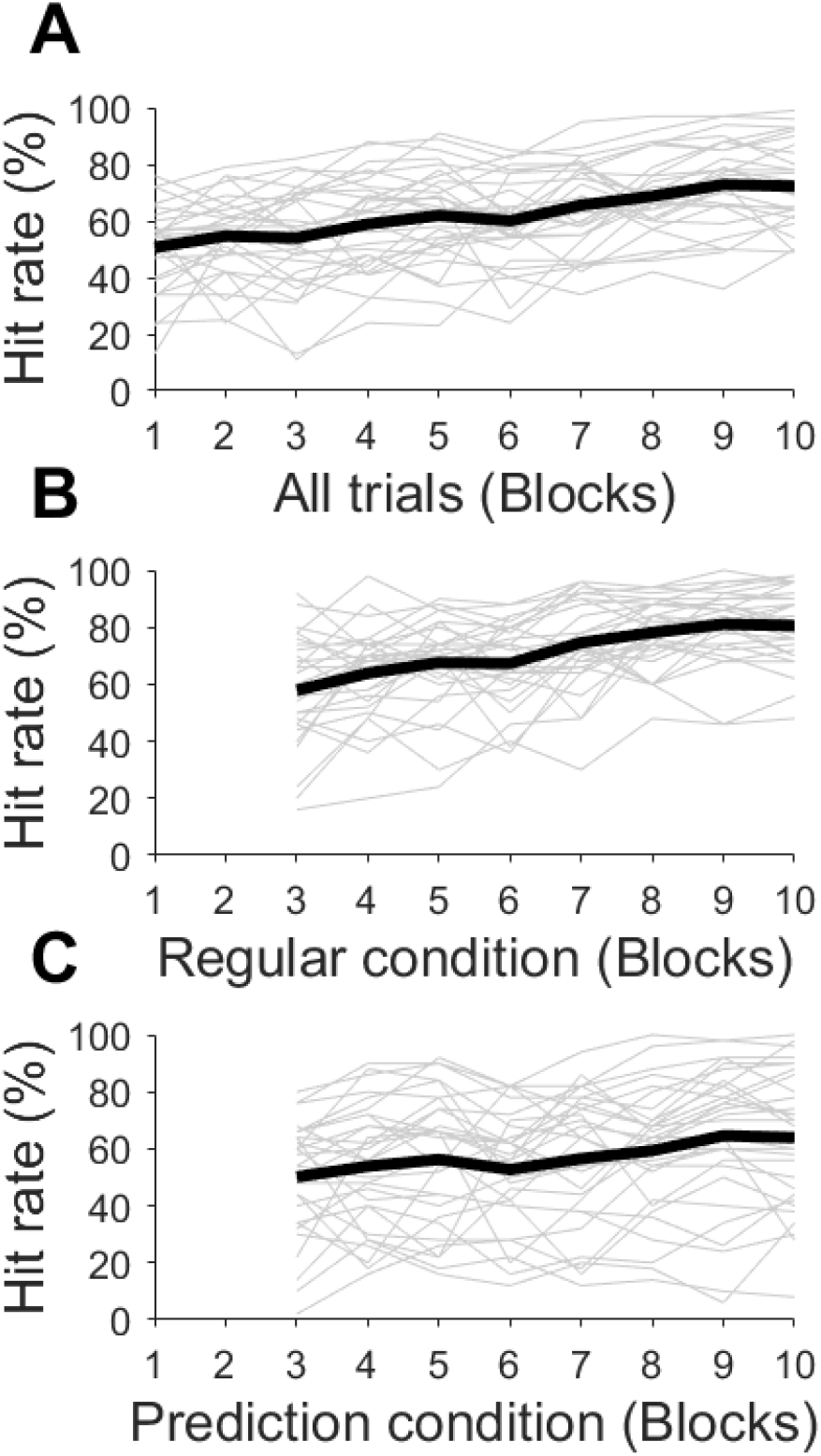
Development of throwing performance over all trials (A), over trials under the regular condition (B), and over trials under the prediction condition (C). Thick black lines represent the group average; thin grey lines represent individual data.

**Table 2.** Post-hoc comparisons between blocks 1-9 and block 10.

| Session | Hit rate<br>Mean difference | SE | t | p<sub>holm</sub> | BF<sub>10</sub> |
| --- | --- | --- | --- | --- | --- |
| 1 vs. 10 | -0.22 | 0.03 | -8.58 | < .001 | > 150 |
| 2 vs. 10 | -0.18 | 0.03 | -6.98 | < .001 | > 150 |
| 3 vs. 10 | -0.19 | 0.03 | -7.24 | < .001 | > 150 |
| 4 vs. 10 | -0.14 | 0.03 | -5.31 | < .001 | > 150 |
| 5 vs. 10 | -0.11 | 0.03 | -4.10 | < .001 | > 150 |
| 6 vs. 10 | -0.01 | 0.03 | -4.85 | < .001 | > 150 |
| 7 vs. 10 | -0.13 | 0.03 | -2.66 | .15 | 6.094 |
| 8 vs. 10 | -0.07 | 0.03 | -1.40 | 1.00 | 1.760 |
| 9 vs. 10 | -0.04 | 0.03 | 0.23 | 1.00 | 0.210 |

**Table 3.** Descriptive data of hit rates in the two experimental conditions and both conditions together (All trials)

|  | N | Mean | SD |
| --- | --- | --- | --- |
| Hit rate *All trials* | 30 | 64.28 | 13.49 |
| Hit rate *Regular* | 30 | 72.26 | 13.77 |
| Hit rate *Prediction* | 30 | 56.00 | 22.71 |
| Difference between<br>*Prediction and Regular* | 30 | 16.27 | 20.73 |

*SD = Standard deviation*

### 3.2 Prediction accuracy

The prediction baseline was on average 55.41 % (SD = 12.83 %), which corresponds to the average chance level for the participants’ predictions. Average prediction accuracy (%ACC_Pred_) exceeded the prediction baseline by 6.01 % (SD = 9.61 %) of the potential accuracy gain by predicting, which was significant (*t*(29) = 3.42, *p* = .002, *d* = .63, *BF*_*10*_ = 19.19). Prediction accuracy, however, varied greatly between participants (see Fig. 3). There was a significant positive correlation of %Acc_Pred_ and hit rate over all trials (*r* = .44, *p* = .014, *BF*_*10*_ = 4.03; Fig. 3A). Since throwing performance differed between the *regular condition* and the *prediction condition* with large variance between participants, it was tested whether this variance also accounted for the differences in prediction accuracies. A significant negative correlation of %Acc_Pred_ with the difference in hit rate between the *regular condition* and the *prediction condition* was found (*r* = -0.52, *p* = .003, *BF*_*10*_ = 4.62; see Fig. 3B). This means that participants with lower hit rates in the *prediction condition* relative to the *regular condition* showed poorer prediction accuracy, in the lowest cases even below baseline level.

**Fig. 3.**
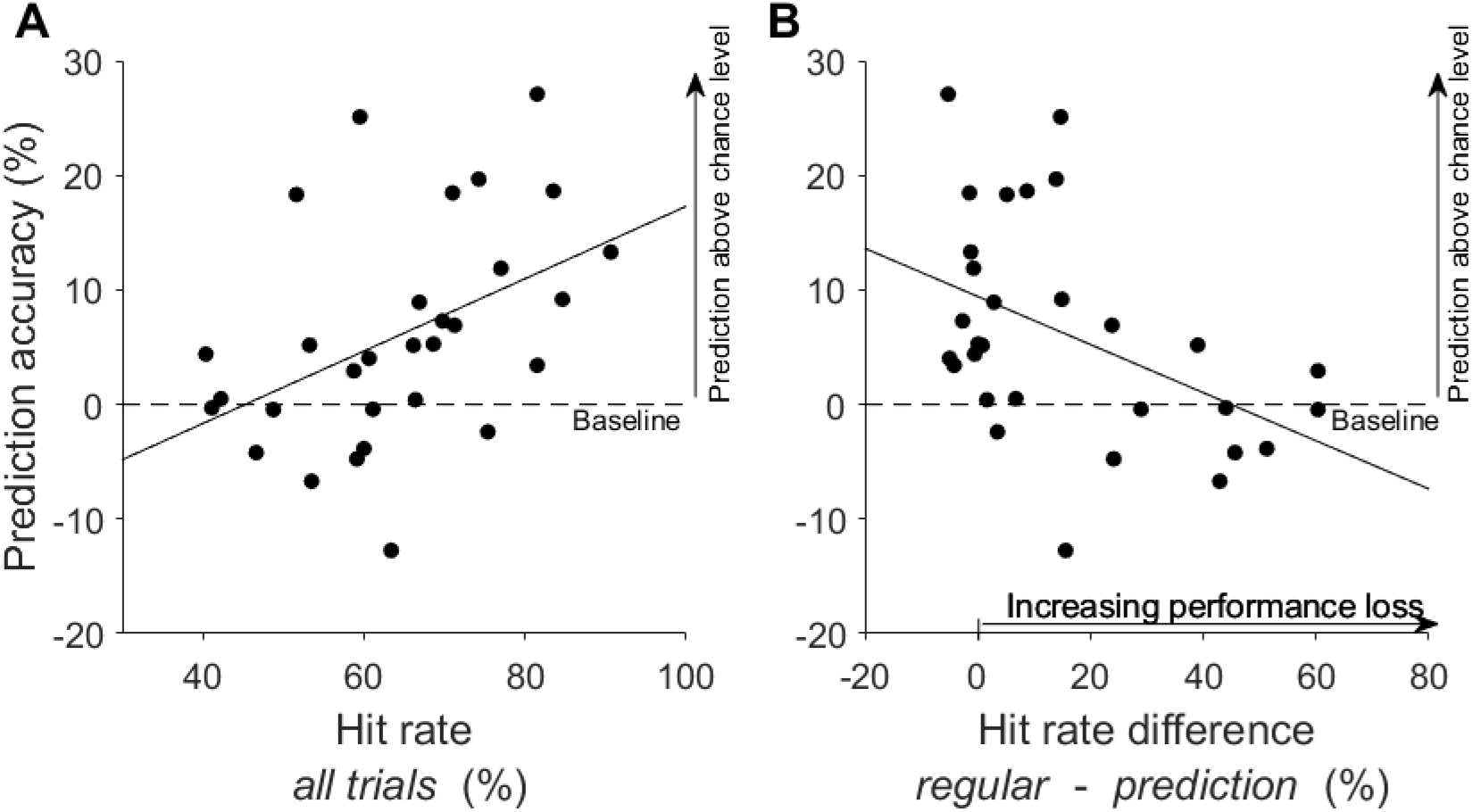
Correlations of throwing performance with prediction accuracy relative to individual chance level (see 2.6 **“**Analysis of verbal responses”). Each dot represents the average values of a single participant. The dashed line marks the baseline prediction level (equivalent to chance level). **A:** Correlation of prediction accuracy with hit rates over all experimental trials (*prediction condition* and *regular condition* together). **B:** Correlation of prediction accuracy with the difference in hit rates between the *prediction condition* and the *regular condition*. The larger the difference value, the lower the performance in the *prediction condition* .

### 3.3 Speech characteristics

Figure 4 shows the response latencies (A) and amplitudes (B) with respect to the *verbalized prediction as a function of actual outcome*. It can be observed that responses predicting misses were generally slower, and tended to also be quieter, than responses predicting hits. The latency difference was confirmed by a main effect regarding the *verbalized result* (*F*(1, 28) = 12.90, *p* = .001, *η*_*p*_^*2*^ = .32, *BF*_10_ > 150), but there was no main effect of *verbalized result* for the amplitude variable (*F*(1, 28) = 0.79, *p* = .38, *η*_*p*_^*2*^ = .03, *BF*_*10*_ *=* .26). There was also a small main effect for *actual result* in response latencies (*F*(1, 28) = 4.78, *p* = .037, *η*_*p*_^*2*^ = .15), which could, however, be ascribed to an interaction effect (see below). Bayesian statistics confirmed that the data was not sufficiently informative to allow a strong conclusion to be drawn about the main effect *actual result* (*BF*_*10*_ = 0.434). Nevertheless, later responses of trials being incorrectly predicted as misses could clearly be observed relative to trials correctly predicted as misses, and this difference was not observable in the trials predicted as hits. Classical ANOVA revealed an interaction effect between *actual result* and *verbalized result* (*F*(1, 28) = 5.41, *p* = .028, *η*_*p*_^*2*^ = .16). A Bayesian mixed-factor ANOVA also determined that the data were well represented by a model that included both main factors, *actual result* and *verbalized result*, and the *actual* × *verbalized* interaction. The *BF*_*10*_ was 7459, indicating decisive evidence in favor of this model when compared to the null model. Moreover, the *BF*_*10*_ in favor of indicating the interaction effect on top of the main effect *actual result* was 5.82. A tendency for a similar interaction (incorrectly predicted misses seem to be expressed most quietly) could be observed in amplitude data, which was, however, neither confirmed by classical nor by Bayesian ANOVA (*F*(1, 28) = .57, *p* = .46, *η*_*p*_^*2*^ = .02, *BF* = .01). There were also no interactions with the covariate *performance loss* (differences in hit rates between the *regular condition* and the *prediction condition*) in the amplitude results.

**Fig. 4.**
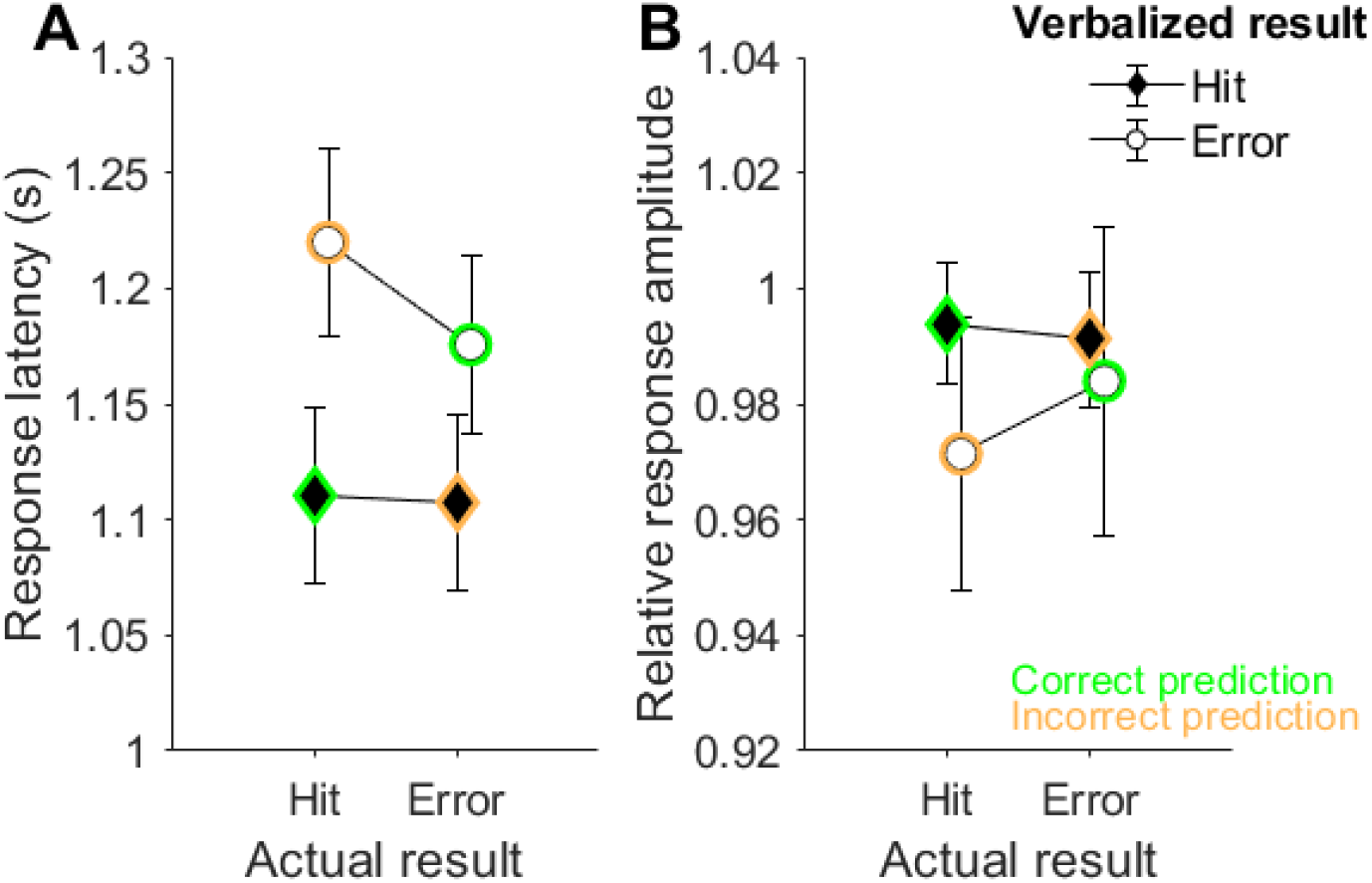
Average response latencies (**A**) and response amplitudes (**B**) differentiated after actual results of trials (actual hits or misses) and the verbalized responses (verbalized hits or misses). Error bars represent standard errors of the mean. Correction predictions are marked in green, incorrect predictions are marked in orange.

Regarding response latency, the three-way interaction with the covariate *actual result* × *prediction* × *performance loss* did not achieve significance, but showed a trend for a different interaction depending on the size of performance loss (*F*(1, 28) = 5.50, *p* = .07, *η*_*p*_^*2*^ = .11). In addition, the Bayes model, including the main factors *actual result* and *verbalized result* and the *performance loss* covariate, strongly outperformed the null model (*BF*_*10*_> 150), and weakly outperformed the model including the *actual* × *verbalized* interaction (*BF*_*10*_= 1.48). To analyze how this trend was supported by the data, a median split was applied with respect to *performance loss*: one group showed virtually no performance loss (< 7.76 %) and the other showed performance loss (> 7.76 %). Figure 5 illustrates that the interaction between *actual result* and *prediction* exists only in the group of participants who showed no performance loss in the *prediction condition*.

**Fig. 5.**
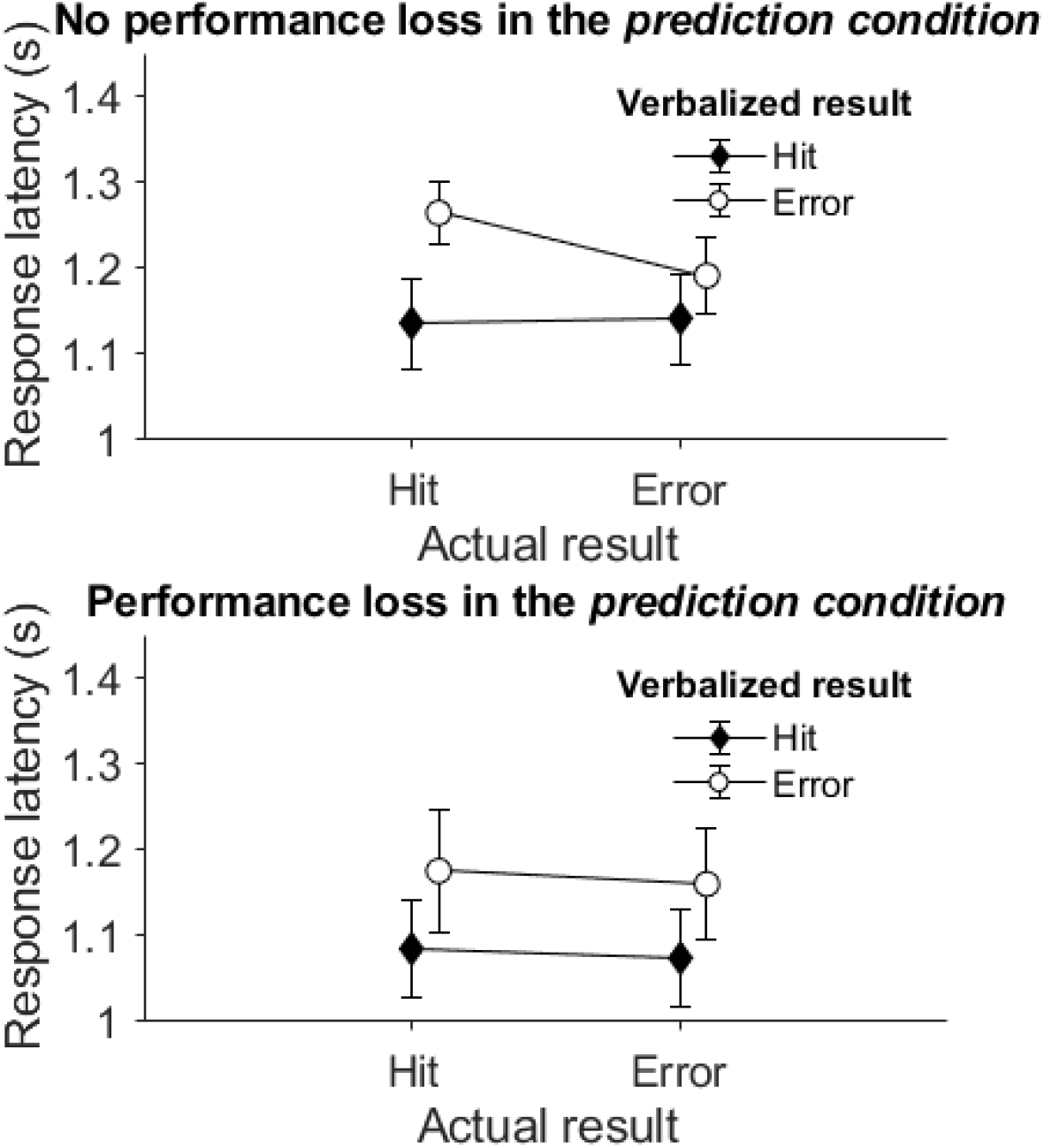
Average response latencies differentiated after actual results of trials and verbalized results for two groups separated based on whether they experienced performance loss in the *prediction condition* or not. Error bars represent standard errors of the mean.

## 4 Discussion

The present study examined whether subjects experienced in a motor task can consciously predict outcomes of their own actions without receiving any external feedback about action outcomes. To this end, participants practiced a virtual goal-oriented throwing task for 200 trials. In every other of the remaining trials, they were then asked to verbally predict whether the ball they had just released would hit or miss the target (*prediction condition*). They had 2.5 seconds after releasing the ball to make their predictions. No feedback was given about the trajectory of the ball. In the other half of the trials, participants did not have to predict outcomes and they could see the ball moving towards the target and hitting or missing it (*regular condition*). Results in terms of throwing performance, prediction accuracy, and speech characteristics of verbal responses were analyzed and compared between the *prediction condition* and the *regular condition*.

### Conscious access to outcome predictions is possible, but varies interindividually

On average, prediction accuracy (as the measure of conscious access to outcome predictions) exceeded baseline levels by about 6 % of the potential gain in accuracy. This means that 6 % of what, in theory, could have been achieved above chance when using internal information was achieved. 100 % in that measure would indicate predictions where every trial is correctly classified as error or hit, while 0 % represented the prediction accuracy that can be achieved without conscious access to internal prediction processes. In this case, verbalized reports of anticipated outcome predictions could only be made at the chance level. Chance level was reflected in the individual baseline level, taking into account actual hit rates (%Act_Hit_ and %Act_Miss_) and verbal report rates (%Verb_Hit_ and %Verb_Miss_). Thus, for accuracies above this level, any improvement in accuracy must have resulted from information gathered during the throwing movement, particularly from internal sources including correlates of efferent commands. Variance of above-baseline gains in accuracy prediction was relatively large between participants, with some individuals achieving gains of around and above 20 %, while others fluctuated around chance level. As described in the introduction, predictive accuracy is a function of prediction quality (quality of a task forward model) and ease or efficiency of conscious access to the forward model predictions. So, individuals with higher prediction accuracies must have had good prediction quality and easy access to their internal processes, while the reason for poorer prediction accuracy may have been poorer prediction quality and/or more difficult access to internal processes, as quality of prediction has already been associated with experience or expertise. Forward model computations contribute to learning (Jordan & Rumelhart, 1992), and extended experience in a motor task correlates with distinct signs of predictive error processing on the neurophysiological level (Lutz et al., 2013; Beaulieu et al, 2014; Maurer et al. 2015; Joch et al., 2017; Maurer et al., 2021). Furthermore, it has been shown that motor experts (sports athletes) can anticipate the outcomes of other players’ actions with relatively little information about action parameters, and that this ability rises with skill level (Abreu et al., 2012; Aglioti et al., 2008; Li & Feng, 2020; Tomeo et al., 2013). In these studies, temporal occlusion paradigms were used, where participants with varying expertise levels watched videos of motor actions and had to predict the outcomes of the actions shown at different points in time. In the Aglioti and colleagues’ study (2008), professional basketball players were capable of correctly predicting shooting outcomes above the level of chance even before players on the videos released the ball. These motor experts, which professional players are, were contrasted with participants with high visual experience (sports journalists and basketball coaches). Pure visual experts needed significantly more information to correctly judge the outcomes above the level of chance. This difference indicates that motor expertise facilitates perceptual abilities in general and, more specifically, at least partially through predictive functions.

The present study extended these findings to predictive abilities with respect to subjects’ own movement outcomes. Although experience and expertise in the task used here was not comparably high to natural experts in terms of practice trials, participants reached a learning plateau, and similar studies with the same task have shown that hit rates of more than 60 % coincided with a neurophysiological marker of forward model predictions (Maurer et al., 2015, Joch et al., 2017). In the present study, participants reached an average hit rate of over 50 % within the first 100 trials, and increased their hit rate average to 72 % in the last 100 trials, taking into account both experimental conditions (*prediction* and *regular*). In the *regular condition* alone, they even reached a hit rate of over 80 %, although interindividual variance in performance was relatively high. Thus, individuals with higher hit rates could be assumed to have had a better forward model of the task and, hence, better prediction quality than individuals with lower hit rates. This was demonstrated by a positive correlation between prediction accuracy and hit rates. The second factor of prediction accuracy, ease of access, also contributed to the clarification of variances between subjects. There was a clear difference in hit rates between the *regular condition* and the *prediction condition*. Hit rates in the *regular condition* were on average higher, but there were again large differences between participants. While about half of the participants did not show much change in hit rates between the two conditions, the other half experienced large decreases in performance when throwing outcomes had to be predicted verbally. This difference in hit rates between the *regular condition* and the *prediction condition* correlated negatively with prediction accuracy. Prediction accuracy was higher when performance loss in the *prediction condition* was smaller. This means that the experimental demands of the *prediction condition* affected motor task performance more in some individuals than in others. What may be the reason for this? During the experimental procedure, participants had to respond verbally no later than 2.5 seconds after releasing the ball. Hence, preparation of the response and the focus on accessing relevant internal information for the outcome prediction might have interfered with motor control processes, leading to a performance loss in the *prediction condition*. This observation is in line with well-established findings that have shown that attention to performance can become counterproductive (Masters, 1992; Masters et al., 1993). This detrimental effect is suggested to be strongest with skill-focused attention (internal focus) of step-by-step monitoring and control (Beilock et al., 2002; Beilock & Carr, 2001; Wulf et al., 1998). Conscious access of internal prediction processes, however, does not necessarily require attention to step-by-step components of a movement. The activation of this access may be more comparable to an external, goal-oriented focus of attention, which typically only affects less-skilled participants (Wulf & Su, 2007). Thus, the less-skilled participants in the present study were more impaired by the verbal prediction requirement, presumably because they had less effective access to information relevant for outcome predictions, and needed to reallocate attention from motor execution. The results from analysis of the speech characteristics of the verbal responses support this interpretation.

### Response latency and amplitude are related to throwing performance and prediction accuracy

Speech production is sensitive, and hesitation and uncertainty of responses can be observed in measures of response latency and amplitude (Seymour, 1970; Collins et al., 2000). Hence, possible detrimental effects of conscious access to movement outcome predictions may be represented by these variables. Movement outcome predictions are based on different sources of information gathered during movement planning (efference copy) and movement execution (haptic, proprioceptive, or visual information; Wolpert et al., 1995), with each of these modalities producing different time delays and resulting in varying degrees of accuracy (Cameron et al., 2014; Pasma et al., 2015; Thorpe et al., 1996). It can be assumed that outcome estimates are continuously produced, resulting in an increase in accuracy of the input information while the movement is evolving. Hence, responses can be quick if sufficiently accurate information is available early on, or if predictions are, instead, based on experience, or are “thoughtlessly” uttered without the subject’s consideration of the actual input information when making predictions.

In the present study, there was a general observation of quicker (and in tendency also louder) responses when a hit was predicted, irrespective of whether this turned out to be true or not. In contrast, response latency differed between correctly and incorrectly predicted misses: Predictions of trials that were incorrectly predicted as misses were verbalized more slowly than predictions of trials correctly verbalized as misses. This difference was only observable in participants without performance loss in the *prediction condition*. Moreover, correctly predicted hits and correctly predicted misses had similar response latencies (and amplitudes), especially in the “no performance loss” group. Hence, in these cases, input information was probably clear and accurate relatively early, leading to relatively fast outcome estimates. In contrast, incorrectly predicted misses resulted in slower response latencies. This indicates that the information gathered here was more ambiguous for outcome predictions, and that this ambiguity did not resolve over time. Nevertheless, participants waited longer, apparently hoping for better information resolution, and ended up answering incorrectly. That this effect was not observed in the “performance loss” group indicates limited access to internal prediction processes in these participants. They were apparently not able to differentiate between accurate and ambiguous input information and, hence, showed similar response times between correctly and incorrectly predicted trials. As already described, the latency effect was generally not present in incorrectly predicted hit trials. The reason for the difference between hit and miss predictions could be a general bias toward hit responses, as has been observed in other studies (Canal-Bruland et al., 2015; Maglott et al., 2019). Response bias in the computation of prediction accuracy was controlled for, but it may have still been inherent in verbal responses. Hence, it is possible that responses predicting hits were expressed based on experiences instead of waiting for accurate input information, which led participants to respond (too) early. However, further experiments would be needed to confirm this assumption.

Taking together results of prediction accuracy, throwing performance, and speech characteristics of verbal responses, it can be concluded that task expertise allows rapid access to accurate motor predictions without interference with motor control processes. On the contrary, when throwing performance is poor, conscious access to internal predictions negatively affects movement execution, presumably in the form of conscious processing costs. In light of these results, the question posed in the title can be answered in the affirmative: Yes, he can! Given the fact that Stephen Curry has exceptionally good throwing skills, it can be assumed that he has an excellent forward model and good access to internal predictions.

### Limitations

There are limitations in this study that need to be taken into consideration when interpreting results. First, no neurophysiological measures were recorded, which could have provided more direct information about internal prediction processes. Temporal characteristics of error-related potentials in electroencephalogram measures, such as the error-related negativity (Falkenstein et al., 1991; Gehring et al., 1993) could have supported interpretations about the response latency effects. In addition, there might be another explanation for the variances in prediction accuracy aside from conscious processing costs that cannot be fully ruled out. In the prediction condition, participants had no visual feedback about throwing effects (i.e., the ball was masked as soon as it was released). Although feedback in the form of information about ball trajectory could not have had any influence on performance in the current trial (since the throwing movement was already terminated at that point), such feedback might have affected the participants anticipatively. That is, knowing that a trial would be without feedback could have unsettled and blocked them. This explanation is regarded as not very likely, but only an experimental separation of missing visual feedback information about action outcomes from conscious processing costs can provide a clear differentiation.

## 5 Acknowledgments

This research was funded by the Deutsche Forschungsgemeinschaft (DFG, German Research Foundation) – project number 222641018 – SFB/TRR 135 TP B6.

